# What can Ribo-seq and proteomics tell us about the non-canonical proteome?

**DOI:** 10.1101/2023.05.16.541049

**Authors:** John R. Prensner, Jennifer G. Abelin, Leron W. Kok, Karl R. Clauser, Jonathan M. Mudge, Jorge Ruiz-Orera, Michal Bassani-Sternberg, Eric W. Deutsch, Sebastiaan van Heesch

**Affiliations:** Department of Pediatrics, Division of Pediatric Hematology/Oncology, University of Michigan Medical School, Ann Arbor, MI 48109, USA; Broad Institute of MIT and Harvard, Cambridge, MA, 02142, USA; Princess Máxima Center for Pediatric Oncology, Heidelberglaan 25, 3584 CS, Utrecht, the Netherlands; European Molecular Biology Laboratory, European Bioinformatics Institute, Wellcome Genome Campus, Hinxton, Cambridge CB10 1SD, UK; Cardiovascular and Metabolic Sciences, Max Delbrück Center for Molecular Medicine in the Helmholtz Association (MDC), 13125 Berlin, Germany; Ludwig Institute for Cancer Research, University of Lausanne, Agora Center Bugnon 25A, 1005 Lausanne, Switzerland; Department of Oncology, Centre hospitalier universitaire vaudois (CHUV), Rue du Bugnon 46, 1005 Lausanne, Switzerland; Agora Cancer Research Centre, 1011 Lausanne, Switzerland; Institute for Systems Biology (ISB), Seattle, Washington 98109, USA

**Keywords:** Ribo-seq, mass spectrometry, immunopeptidomics, non-canonical open reading frame, microprotein

## Abstract

Ribosome profiling (Ribo-seq) has proven transformative for our understanding of the human genome and proteome by illuminating thousands of non-canonical sites of ribosome translation outside of the currently annotated coding sequences (CDSs). A conservative estimate suggests that at least 7,000 non-canonical open reading frames (ORFs) are translated, which, at first glance, has the potential to expand the number of human protein-coding sequences by 30%, from ∼19,500 annotated CDSs to over 26,000. Yet, additional scrutiny of these ORFs has raised numerous questions about what fraction of them truly produce a protein product and what fraction of those can be understood as proteins according to conventional understanding of the term. Adding further complication is the fact that published estimates of non-canonical ORFs vary widely by around 30-fold, from several thousand to several hundred thousand. The summation of this research has left the genomics and proteomics communities both excited by the prospect of new coding regions in the human genome, but searching for guidance on how to proceed. Here, we discuss the current state of non-canonical ORF research, databases, and interpretation, focusing on how to assess whether a given ORF can be said to be “protein-coding”.

**In brief:** The human genome encodes thousands of non-canonical open reading frames (ORFs) in addition to protein-coding genes. As a nascent field, many questions remain regarding non-canonical ORFs. How many exist? Do they encode proteins? What level of evidence is needed for their verification? Central to these debates has been the advent of ribosome profiling (Ribo-seq) as a method to discern genome-wide ribosome occupancy, and immunopeptidomics as a method to detect peptides that are processed and presented by MHC molecules and not observed in traditional proteomics experiments. This article provides a synthesis of the current state of non-canonical ORF research and proposes standards for their future investigation and reporting.

**Highlights:**

- Combined use of Ribo-seq and proteomics-based methods enables optimal confidence in detecting non-canonical ORFs and their protein products.
- Ribo-seq can provide more sensitive detection of non-canonical ORFs, but data quality and analytical pipelines will impact results.
- Non-canonical ORF catalogs are diverse and span both high-stringency and low-stringency ORF nominations.
- A framework for standardized non-canonical ORF evidence will advance the research field.

**Graphical Abstract:** 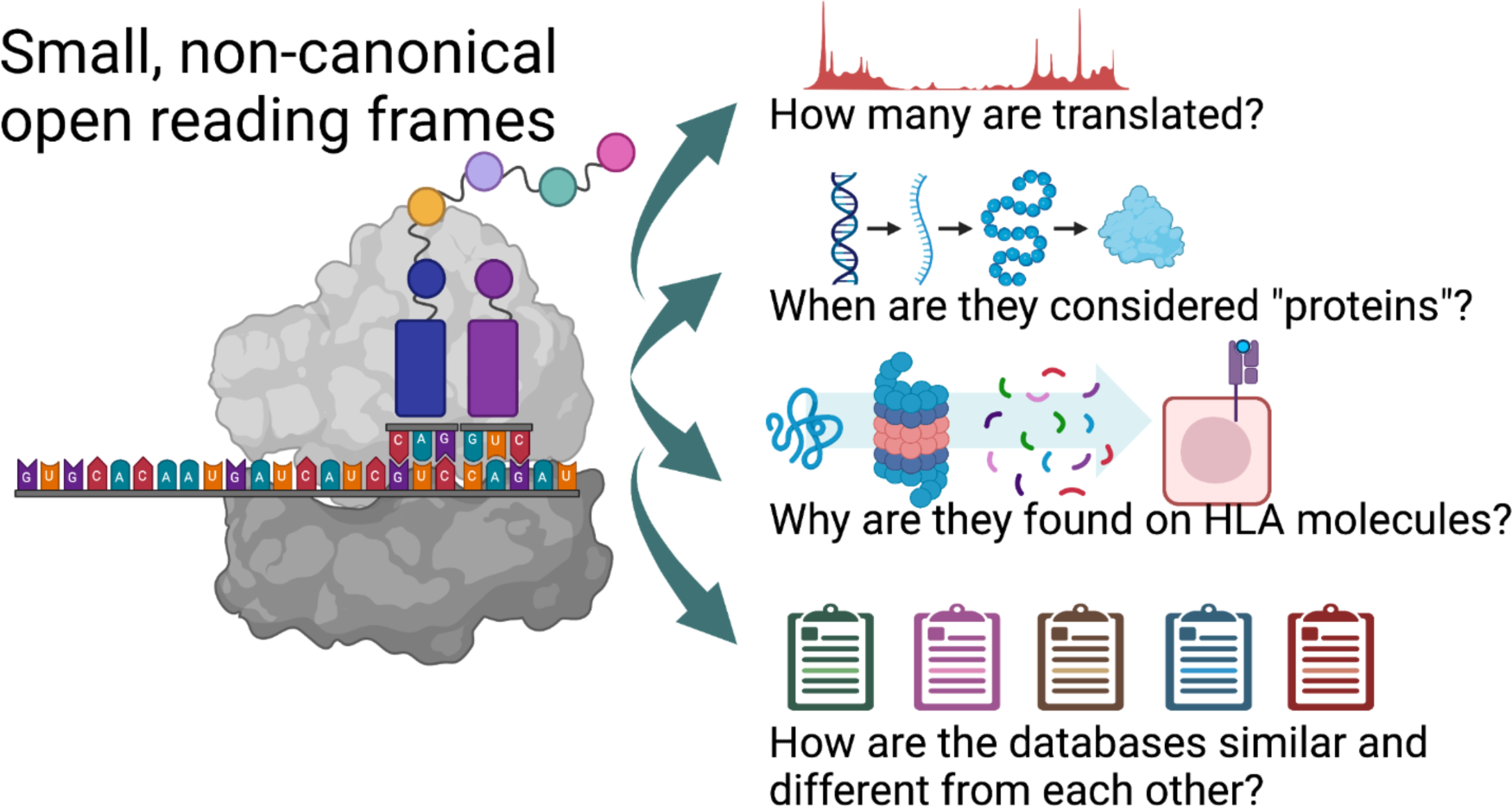

## Introduction

Defining the extent of RNA translation in the human genome – and the resulting proteins – has long been a major focus for biomedical research. Approximately 19,500 protein-coding genes, which produce ∼80,000 annotated protein coding isoforms, constitute the canonical proteome (1–6). Yet, whether this catalog is comprehensive has recently undergone substantial debate spurred by sequencing-based advances in the analysis of ribosome translation, termed ribosome profiling (Ribo-seq). Based on classical techniques used to isolate ribosome-RNA complexes, Ribo-seq is a RNA sequencing-based approach that profiles ribosome-protected RNA fragments, precisely defining open reading frames (ORFs) actively engaged by translating ribosomes (7, 8). As a tool to detect the translation of RNA, the precision of this methodology is unprecedented: from individual ribosome footprints, the exact codon being translated in a purified ribosome-RNA complex can be determined. Through the sequencing of hundreds of millions of ribosome footprints, a single Ribo-seq experiment can therefore produce a detailed and accurate representation of a given sample’s translated RNAs, typically identifying ∼11,000 - 12,000 translated genes per sample (9–11), which is more similar to the ∼12,000 - 13,000 expressed protein-coding mRNAs detected in a given cell type (12) compared to the ∼9,000 - 11,000 proteins per sample typically detected in mass spectrometry methods (13, 14).

In addition to confirming known protein-coding sequences (CDSs), the high predictive power of Ribo-seq has unveiled thousands of other genomic sites of ribosome translation. These are most commonly found within known mRNAs (i.e., different reading frames than canonical CDS regions), but also within transcripts annotated as long noncoding RNAs, pseudogenes, or retroviral elements in the genome (7, 9, 11, 15–23). Ribo-seq can also provide clues on previously missed N-terminal in-frame extensions to known CDSs, initiated at sites alternative to the classically annotated initiation codon (24–27). The nomenclature and estimated abundance of non-canonical ORFs are listed in **Figure 1A**. For clarity, these ORFs are termed “non-canonical” to distinguish them from CDSs included in reference gene annotation - i.e. Ensembl-GENCODE - even though their translation, to our knowledge, occurs through mechanisms of ribosome activity similar to that of CDSs. Throughout this text the term “non-canonical open reading frame” is therefore defined as any open reading frame that is not an annotated CDS, an in-frame extension or truncation (either N-terminal or C-terminal), or an in-frame intron retention of an annotated CDS. For our purposes, we will be focusing on upstream ORFs (uORFs), upstream overlapping ORFs (uoORFs), internal ORFs that overlap the CDS but are translated in a different frame (intORFs), downstream overlapping ORFs (doORFs), downstream ORFs (dORFs), and lncRNA-ORFs (as in **Figure 1A**). We will not discuss in depth ORFs that may be translated from pseudogenes (19), genomic retroviruses (28), or other repetitive sequences (29) (See **Limitations** section).

**Figure 1:**
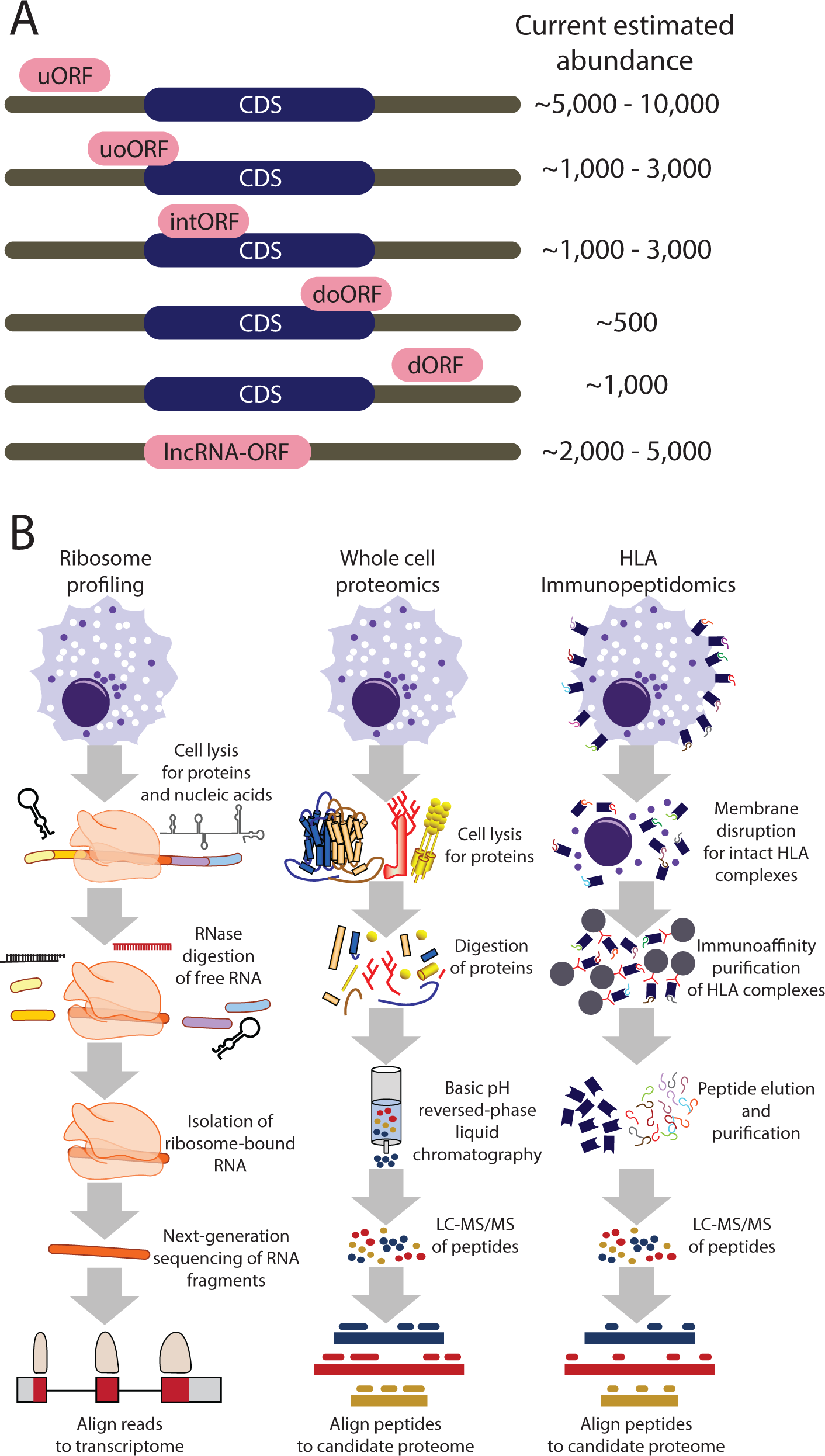
An overview of non-canonical ORF types and detection methods. A) A schematic illustrating the standardized names of non-canonical ORF types, their relationship to known mRNAs, and current estimations of their abundance. CDS, protein-coding sequence; uORF, upstream open reading frame (ORF); uoORF, upstream overlapping ORF; intORF, internal ORF; doORF, downstream overlapping ORF; dORF, downstream ORF; lncRNA-ORF, ORF residing within an annotated lncRNA. B) Generalized workflows for ribosome profiling (Ribo-seq), tryptic whole cell mass spectrometry, and HLA immunopeptidomics. The schematic indicates general properties of sample preparation for these data types.

Given these observations, the genomics community has been faced with the fundamental question: does the genome actually encode far more than the ∼19,500 protein-coding genes currently accepted as canonical? In response, there have been increasing efforts to corroborate the observations from Ribo-seq using mass spectrometry, with the overall conclusion that only a low percentage of non-canonical ORFs are detectable by conventional tryptic whole proteome methods employing liquid chromatography with tandem mass spectrometry (LC-MS/MS) techniques (9, 15, 30–34). Yet, far more non-canonical ORFs appear to be detectable with immunopeptidomic approaches that profile peptides presented by the class I human leukocyte antigen (HLA-I) system (**Figure 1B**) (34–39). Moreover, independent of their protein-coding capacity, non-canonical ORFs may serve important roles in the regulation of mRNA translation (40–42). With these observations at hand, one of the central tasks for the proteomics and genomics communities alike is to develop a consensus understanding on what constitutes sufficient evidence of detection for a non-canonical ORF from each technology and how to standardize these assessments given the limitations of each methodology.

### Types of evidence for non-canonical ORFs

Translated non-canonical ORFs can be detected by either Ribo-seq or LC-MS/MS approaches, with examples of transition to canonical annotated protein-coding genes emerging from both. For example, translation of the signaling proteins APELA (43), POLGARF (44, 45), TINCR (46) and the cardiac proteins MYMX (47) and MRLN (48) was first identified using Ribo-seq, while shotgun LC-MS/MS data provided the initial evidence for the translation products of uORFs in ASNSD1, MKKS, MIEF1, and SLC35A4 (30, 49).

Together, the combination of Ribo-seq and LC-MS/MS is a powerful way to identify translated CDSs and ORFs (21, 50–52). Ribo-seq does not directly detect proteins, but rather provides evidence of ongoing nucleotide translation. By contrast, LC-MS/MS evidence for non-canonical ORFs takes the form of direct detection of peptides. In the case of conventional LC-MS/MS of cellular lysates, these peptides are typically tryptic, meaning they were generated by protein cleavage at the C-terminal side of a lysine or arginine, or semi-tryptic, meaning they were generated by protein cleavage at the C-terminal side of a lysine or arginine at one end of the peptide but not the other. However, many ORFs have now been observed in mass spectrometry-based HLA-I immunopeptidomics data (18, 34, 36, 38, 53). Here, no tryptic digestion is employed. Instead, peptides containing the HLA-I peptide binding motifs of the HLA-I allele expressed by a specific cell line or tissue are observed. A variety of lower-throughput approaches have also been used to assess translation of non-canonical ORFs, including generation of custom antibodies, expression of epitope-tagged ORF cDNAs, selective reaction monitoring (SRM), and radiolabeled *in vitro* translation (9, 17, 54–56).

While high-quality Ribo-seq and LC-MS/MS whole proteome data on the same sample should be able to identify highly consistent sets of endogenous CDSs, Ribo-seq is not able to pinpoint the responsible translation event for exogenous proteins, which originate from sources other than the sample’s own genetic material. Similarly, Ribo-seq cannot detect or predict protein stability, folding, or post-translational modification. If there is a substantial discrepancy with MS detecting many additional proteins, then the quality of the Ribo-seq library should be inspected (see below). It should also be noted that Ribo-seq, like all sequencing-based methods, may not be able to resolve translation events in repetitive genomic regions, such as retrotransposons, pseudogenes, or genes with very high homology.

By contrast, Ribo-seq will almost always detect many non-canonical ORFs that are not found by proteomics. This is due to several factors: both the nature of the data itself as well as technological differences in the methods that may impact the ability to detect lowly-expressed molecules with high confidence. For example, all mass spectrometry-based proteomics methods lack a PCR amplification step that is present in most sequencing-based methods, which enables higher sensitivity at lower sample inputs. Regarding the nature of the data, Ribo-seq has the ability to identify translating ribosome signatures in an unbiased way, which may confidently find ORFs less than 8 amino acids long that are fundamentally challenging to identify by MS (15, 57). In fact, Ribo-seq can confidently identify an ORF that is simply a start codon followed by a stop codon (i.e. Met*), because the Ribo-seq reads remain sufficiently long for unique genomic mapping (58).

Second, since some non-canonical ORFs are located in GC-rich promoters (such as uORFs), these may encode amino acid sequences that are enriched in arginine (CGU/CGC/CGA/CGG codons) and thus would be excessively cleaved by trypsin to small peptides that cannot be uniquely mapped to a single ORF. Whether use of alternative proteases (59) could improve non-canonical ORF detection in whole lysate proteomics is unclear.

### Considerations and quality control steps for the data-driven discovery of non-canonical human ORFs

Differences in the nature of Ribo-seq and LC-MS/MS-based whole proteome and immunopeptidome data collection also represent a source of substantial variability in the detection of non-canonical ORFs. Notably, while targeted whole proteome and immunopeptidome LC-MS/MS approaches may offer improved sensitivity, these require candidate non-canonical proteins of interest to be known prior to analysis. While each method uses high-throughput data generation to profile cellular translation comprehensively, the data have intrinsically different strengths and weaknesses that may result in discordance between them (**Table 1**).

**Table 1.** Features and characteristics of methods to detect non-canonical ORF translation.

|  | Molecule detected | Digestion step? | Target size of analyte | Number of annotated CDSs detected | Number of ORFs detected | Strengths | Weaknesses |
| --- | --- | --- | --- | --- | --- | --- | --- |
| Ribo-seq | RNA bound within ribosomes | RNase and DNase | 28 - 30 nt | 10,000 - 13,000 | 2,000 - 200,000 | -Genome-wide<br>-No bias due to trypsin<br>-Detects small and large CDSs<br>-Nucleotide-level precision<br>-Defines exact reading frame of ORF | -Does not detect proteins directly<br>-Cannot detect PTMs<br>-Cannot inform post-translational protein regulation<br>-Analysis pipelines may be discordant |
| LC-MS/MS | Tryptic peptides | Trypsin | 8 - 25 amino acids | 9,000 - 11,000 | 10s to 100s | -Direct protein detection<br>-Informs protein abundance<br>-May detect PTMs<br>-Proteome-wide | -High false-positive rate without Ribo-seq<br>-Biased against small proteins<br>-Trypsin may bias protein representation<br>-Does not provide nucleotide-level precision |
| HLA Immuno-peptidomics | HLA-presented peptide antigens | None | 8 - 12 amino acids | 8,000 - 10,000 | 1000 - 5000 | -Direct protein detection<br>-Enrichment for low-abundance, strong binders<br>-Proteome-wide<br>-Can detect unstable translations<br>-Does not require tryptic sites | -Does not inform protein stability<br>-Does not indicate intracellular abundance<br>-HLA allele expression limits peptide representation<br>-Does not provide nucleotide-level precision |

#### Ribo-seq

The quality of a Ribo-seq dataset is most commonly evaluated using three considerations: codon periodicity, library complexity, and number of canonical CDSs identified.

Codon periodicity reflects the percentage of Ribo-seq reads that correctly identify the known reading frame of CDSs (**Figure 2A-C**). In a high-quality Ribo-seq dataset, >=70% of reads that are between 28 and 30 nucleotides in length map to the correct reading frame of known CDSs. The precise read length that displays the most preferable (the “cleanest”) signal can vary and depends on the sample type and the method of nuclease digestion used to eliminate cellular RNAs not bound within the translating ribosome. Because of limitations of the experimental technique as well as biological variation in ribosome occupancy, a codon periodicity above 90% is typically not attainable (60). A Ribo-seq dataset with a codon periodicity < 60% should ideally not be used for ORF discovery due to challenges with accurate identification of the reading frame (19, 60, 61). A periodicity between 60 – 70% is a gray zone where the data may be used in some cases with increased caution and stringency.

**Figure 2:**
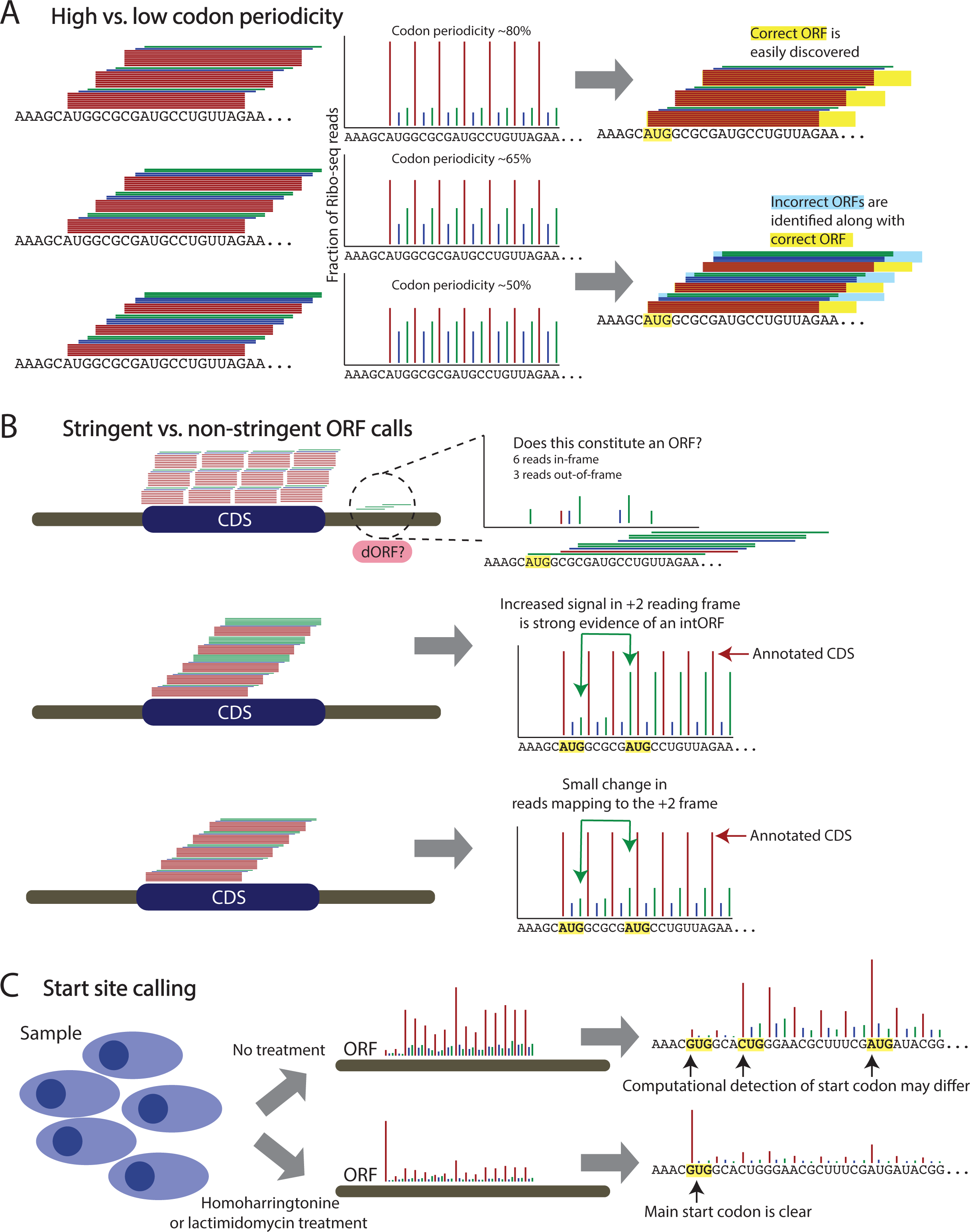
Quality metrics of Ribo-seq and stringency of ORF calling. A) An illustration showing codon periodicity as a central metric of Ribo-seq library generation. Three illustrations indicate high-quality, borderline, and poor-quality Ribo-seq libraries. B) An illustration representing high-stringency and low-stringency ORF calling. In the top case, a small number of reads map the the 3’UTR of an annotated mRNA, and only two-thirds of those 3’UTR reads support the same reading frame of a potential dORF nomination. In the middle and bottom cases, a potential intORF has varying read support evidence. The middle case shows clear evidence of an intORF by a large increase in reads mapping to the +2 reading frame midway through the CDS. In the bottom case, there is a smaller change in the reads mapping to the +2 reading frame. C) Use of ribosome-stalling drug treatments to clarify translational start sites. Cultured cells are treated with homoharringtonine or lactimidomycin to stall ribosomes at the main translational start site of a given ORF, leading to a clearer resolution of the specific start codon.

Library complexity refers to the number of unique RNA molecules sequenced and what fraction of these are ribosome footprints that map to CDSs. The challenge with a low complexity library is that the majority of the reads will be PCR duplicates. When the number of initially isolated footprints is limited (e.g. due to low quality of the input material or sub-optimal sample processing), ultimately many duplicate copies of this limited number of footprints will be sequenced. This means that deeper sequencing of this library will yield no or only minimally more biologically distinct footprints. Typically, the majority of reads in such low complexity libraries will come from non-footprint sources, particularly intergenic and intronic contaminants (e.g. microsatellite repeat elements, ribosomal RNAs, or small RNAs that overlap gene regions), which are unintentionally isolated during the Ribo-seq procedure because these RNA species are of a similar size to the ribosomal footprint and may have certain RNA structures (62, 63). In general, a Ribo-seq library with sufficient complexity will have the majority of reads mapping to annotated and novel CDSs. In some cases, such as with degraded samples, there may be substantial intergenic noise or a higher fraction of RNA species that are normally restricted to the cell nucleus, but yet still sufficient codon periodicity and library complexity in terms of unique RNA molecules that map to CDSs. Here, the challenge is to achieve sufficient sequencing depth to ensure adequate sampling of unique RNA molecules. While 150 million reads typically suffices for the analysis of a high-quality Ribo-seq library, a “noisy” – yet usable – library may require very deep coverage (>400 million reads), which is mostly a consideration for the financial cost of the sequencing (60, 64, 65). For human Ribo-seq libraries, typically 15-30% of the sequenced reads can be classified as ribosome footprints and the rest is often discarded. For a library sequenced to a depth of 150 million reads, that would total to approximately 22.5 to 45 million ribosome footprints – a number comparable to a routinely sequenced RNA-seq library. Of these, >80% should map to annotated CDSs (60), leaving ∼5 million ribosome footprints for ORF discovery.

The number of known CDSs identified is particularly important when one aims to provide a comprehensive view of all translated ORFs in a sample of interest. This metric relates both to the amount of noise in the library, the periodicity of the footprints, as well as the depth of the sequencing. A sufficiently sequenced Ribo-seq library for a human sample with high periodicity should detect at least >9,000 annotated CDSs, and often >10,000 (9–11, 18). Human sample Ribo-seq libraries that do not reach this threshold – despite sufficiently deep sequencing and periodicity – should be used with caution, as the false negative rate for detecting ORFs will be high (many ORFs will be missed). While Ribo-seq-based ORF detection tools theoretically have a low false-negative rate, the confidence (FDR) with which an ORF or CDS is detected, the number of independent samples in which it can be found, and the translation rate of the ORF should always inform research decision-making. For instance, direct comparison of non-canonical ORF FDRs and translation rates, compared to those of canonical CDSs, can inform both the relative abundance of the ORF’s translation product and the degree of certainty with which the algorithm could nominate it.

Because *de novo* and *ab initio* RNA assemblies are technically challenging with the short nucleotide sequences (28 – 30 nt) obtained during a Ribo-seq experiment, analysis of Ribo-seq data requires alignment of the reads to a reference transcriptome, most commonly Ensembl or RefSeq though custom transcriptomes are also used in some cases. Statistical assessment of a non-canonical ORF nomination is inconsistent across computational methods, with some approaches calculating a p-value for significance (e.g., Ribotaper (61), ORFquant (10), Ribo-TISH (66), PRICE (67), RiboCode (68)) and other approaches computing confidence scores (e.g., RibORF (19), Ribotricer (69), ORF-RATER (70)). In addition, these methods are often based on fundamentally different modeling approaches, including Hidden Markov (RiboHMM (20)), multitaper (Ribotaper (61)), transformer (DeebRibo, TIS Transformer (71, 72)), support vector machine (RibORF (19)), expectation-maximization (EM) (PRICE (19, 67)) models, among others. As such, different methods may be more appropriate for certain research questions, datasets or desired ORF types.

As a consequence, two different algorithms can have differing ORF outputs for the same gene. This can be due to the level of stringency or the strengths and weaknesses of a particular ORF caller for a certain type of ORF or certain quality of data. For example, some ORF callers cannot detect ORFs with near cognate start codons, whereas others are better suited for the detection of overlapping reading frames where periodic footprint signals are mixed and hard to dissect. Other tools handle alternative splicing better. Depending on the research question, input data quality, species of interest, or annotation goals, combinations of ORF callers followed by curation of called ORFs may be necessary (see below in **How many non-canonical ORFs are there?**).

#### HLA-I and HLA-II Immunopeptidomics

In the past decade, interest in HLA-I and HLA-II presented peptides has become widespread across many areas of biomedical research, as a subset of HLA-presented peptides demonstrate antigenic properties and represent a class of potential therapeutic targets (73–76). The application of HLA immunopeptidomics differs from tryptic whole proteome protocols, as these methods leverage native lysis buffer and antibody or affinity-tag enrichment steps to isolate HLA-peptide complexes from cell lysates (**Figure 1B**) (77, 78). The peptides are naturally produced following degradation of endogenously expressed source proteins by cellular proteases and peptidases and the proteasome. As such, no tryptic digestion is used in immunopeptidome analyses. Therefore, regarding detection of non-canonical proteins, HLA immunopeptidome analysis has three advantages over tryptic whole proteome analysis: 1) each HLA allele has a distinct peptide-binding motif that presents specific subsets of peptides, which can then be detected with mass spectrometry in the absence of digestion with a protease; 2) the HLA presentation pathway may have privileged access to proteins that are rapidly degraded as the half-life of HLA-peptide complexes (hours) are in general longer than the half-life of rapidly degraded proteins (minutes); and 3) HLA immunopeptidomics broadly samples endogenous proteins from all abundance levels including those from lower-abundance non-canonical ORFs (79–81). These advantages align with recent studies that have shown higher observation rates of non-canonical proteins in the HLA-I immunopeptidome compared to the tryptic whole proteome (39, 82).

Similar to tryptic whole proteome datasets, immunopeptidome datasets require strict quality control steps to ensure the data and analysis are of high-quality. Peptide length, the presence of peptide-binding motifs, and predicted binding to HLA molecules coded by specific alleles are common quality control steps in immunopeptidomics workflows. Because HLA-I and HLA-II molecules have unique peptide binding grooves that accommodate peptides of different lengths, peptide size is an important quality control metric of immunopeptidomics data. Specifically, HLA-I peptides are ∼8-12 amino acids long (mostly 9mers), whereas HLA-II peptides are generally 12-25mers (77). HLA-II peptides are also typically found in nested sets, while this is not a global feature of HLA-I peptides, and can also be used to quality control HLA-II immunopeptidome datasets. Furthermore, each individual person expresses different HLA alleles with distinct HLA-binding motifs, which influence which peptides are presented. Therefore, it is common to confirm that HLA allele-specific binding motifs of the expressed HLA molecules are present in the immunopeptidome data, and that peptides derived from canonical and non-canonical ORFs in a given dataset are predicted to bind to the expressed HLA molecules to a similar extent. A number of computational approaches (e.g. MHCflurry, NetMHCpan, MixMHCpred, ForestMHC, HLAthena) can be used to both predict HLA peptides and the strength of their binding to various HLA molecules (76, 83–88). It is important to note that HLA-I binding prediction is currently more accurate compared to HLA-II binding prediction, as HLA-II motifs are more complex and large subsets of diverse HLA-II heterodimers are in the process of being characterized and the associated prediction algorithms are being further improved (89–92).

Interestingly, peptides derived from non-canonical ORFs are much more abundant in HLA-I datasets compared to HLA-II datasets (18, 34, 36, 38, 39, 53, 93). HLA-I molecules usually present peptides derived from proteasome-mediated degradation of newly synthesized and other cellular proteins, and HLA-I presentation is tightly linked with protein synthesis and degradation rates. In contrast, HLA-II molecules, that are often expressed on professional antigen presenting cells (APCs), present peptides derived from degradation of extracellular proteins that were taken up by the APCs or from endogenous proteins that are destined to be degraded in specialized vacuolar compartments of the endosome-lysosome system. Both HLA-I and HLA-II systems require trafficking to ensure peptide loading in the right compartment. For HLA-I, the peptides themselves are transported into the ER by a transporter associated with antigen processing (TAP), while in case of HLA-II, the source proteins must first reach the acidic compartments for degradation, for example via receptor mediated internalization or recycling of transmembrane proteins. Hence, the sources of HLA–II presented peptides are often stable and abundant proteins.

Because of HLA-I binding constraints, and the short length of some non-canonical proteins, a non-canonical ORF is often represented by a single peptide in HLA-I immunopeptidome data, and therefore additional quality control measures should be taken to support these identifications. To this end, a non-canonical protein subset-specific FDR threshold should be applied to each individual ORF type, rather than a global FDR (82, 94) because non-canonical ORF peptides represent a small fraction (typically <5%) of the overall immunopeptidome and individual ORF types vary considerably in their frequency. Thus, a global FDR can be excessively permissive for a small subpopulation and lead to higher false-positive identifications.

Beyond leveraging known HLA specific peptide lengths, binding motifs and subset-specific FDR, there are further quality metrics that can be applied to immunopeptidomics datasets when the focus is the identification of rare, non-canonical proteins (95). The gold standard for supporting the identification of non-canonical peptides presented by HLA molecules is by comparing the retention time and MS/MS spectrum of an identified peptide with a synthetic peptide of the same amino acid sequence. However, it is often the case that hundreds of non-canonical peptides are identified in a single HLA-I immunopeptidome experiment, making the synthetic peptide confirmation for all potential non-canonical derived HLA-I peptides not feasible. To overcome this challenge, it is now possible to compare the observed MS/MS spectra with predicted MS/MS spectra with tools such as Prosit (96). The comparison of the predicted and observed MS/MS spectra provides additional support for non-canonical peptide identification (97, 98). In addition, there are also multiple algorithms that can predict peptide retention times. The predicted retention time, using tools such as DeepLC or DeepRescore, can be compared to measured retention time for all peptides in a sample (canonical and non-canonical), as the correlation between predicted and observed retention time supports the LC-MS/MS identifications of non-canonical derived peptides in immunopeptidomes (99, 100). Overall, deep learning based prediction of peptide MS/MS spectra and retention time are powerful tools that help reduce the number of false positive non-canonical peptide identifications in immunopeptidome datasets.

#### Tryptic whole proteome LC-MS/MS

Rigorous standards for the analysis of LC-MS/MS tryptic whole proteome data have been established by the Human Proteome Organization/Human Proteome Project (HUPO/HPP) international consortium, as reviewed elsewhere (101–103), and these standards remain the expectation for researchers claiming identification of non-canonical ORF peptides (30). For claims of detection of proteins not previously detected, these guidelines require two non-nested, uniquely mapping peptides each of at least nine residues in length with a total extent of at least 18 amino acids and with high-quality peptide-spectrum matches (PSMs) upon manual inspection (30, 101, 103). These peptide-spectrum matches should be provided in the form of Universal Spectrum Identifiers (USIs) so that the spectra can be easily examined by others (104).

Yet, consistent application of high-quality tryptic whole proteome data collection and analysis guidelines remains non-uniform across the research community. Proteogenomic studies looking for non-canonical ORFs without Ribo-seq data – i.e., by predicting and including all ORFs in RNA transcripts – have been plagued by high false-positive rates (30, 49, 105–108), and initial efforts to inspect early claims of non-canonical ORF peptides concluded that “many of the spectral matches appear suspect” (30).

Moreover, while use of decoys is standard in tryptic whole proteome experiments to define global false discovery rates, decoys may be less useful for distinguishing true peptides for non-canonical ORFs. Indeed, Wacholder *et al.* have concluded that decoy bias among non-canonical ORF products leads to inaccurate FDR estimates for short ORFs when decoys are created by reversing the complete protein sequence, but not when excluding the initial Met from the reversal (109). Lastly, efforts to identify non-canonical ORFs in tryptic whole proteome data must account for peptides instead being derived from canonical variants including single amino acid variants (SAAVs) and splice-site peptides for alternative isoforms of known CDSs. The use of personalized proteogenomic database searches is not straightforward or used by all in the proteomics community.

Considering these factors, the general experience of the research community is that few non-canonical ORFs are found by conventional tryptic whole proteome LC-MS/MS analyses and some of those are ultimately false-positive peptides (110, 111). In some cases, such ORFs are “undiscoverable” by tryptic whole proteome approaches, either due to the short length of non-canonical ORFs or intrinsic sequence features that do not produce LC-MS/MS observable tryptic peptides. For example, translation of repetitive amino acid sequences (e.g. glycine-leucine) has recently been described (29). Nevertheless, even approaches aimed at enriching for small proteins from cell lysates result in only modest increases in non-canonical ORF detection, rather than exponential increases (33). On the other hand, other enrichment techniques focused on post-translational modifications (PTMs; i.e. the acetylome, phosphoproteome, and ubiquitylome) have also reported non-canonical proteins and may provide both an alternative method to enrich for non-canonical proteins and also hint towards potential functional relevance of this subset of non-canonical proteins given the cellular roles of those PTMs (82).

Furthermore, data-independent acquisition mass spectrometry (DIA-MS) provides a potential opportunity to detect non-canonical ORF-derived peptides that have been reliably detected previously with high quality spectra obtained with narrow isolation windows from a data-dependent acquisition (DDA) approach. In DIA-MS, previously identified peptides are more reproducibly sampled by sequentially isolating and fragmenting peptides across the *m/z* range, which decreases stochastic sampling bias toward higher abundant species and may increase the chances of finding rare non-canonical ORFs (112). This approach has been used in conjunction with Ribo-seq to claim detection of microproteins from non-canonical ORFs (50).

Beyond technical limitations of mass spectrometry, there are also biological factors that may make non-canonical ORFs less frequently observed in tryptic whole proteome LC-MS/MS datasets. To this end, there is increasing evidence that points toward intrinsic instability of proteins translated from non-canonical ORFs, resulting in their immediate degradation. Kesner *et al.* used functional genomics approaches to demonstrate that the ribosome-associated BAG6 membrane protein may directly triage hydrophobic non-canonical ORF translations to the proteasome for degradation (113). Thus, it is possible that many non-canonical ORFs do not generate a stable protein product and might only be observable by immunopeptidomics or in whole proteome experiments with inhibition of the protein degradation mechanisms of a cell.

### How many non-canonical human ORFs are there?

The number of non-canonical ORFs encoded in the human genome remains highly speculative. To date, a limited number of human tissues and cell lines have been analyzed by Ribo-seq, and proteogenomics studies that have aimed to incorporate ORFs derived from these datasets have been difficult to interpret due to numerous false positives. As such, while it is well-established that the human genome contains thousands of translated non-canonical ORFs, whether the precise number is closer to 10,000 or 100,000 remains a matter of debate. A further complication is that different research communities may not use a consistent definition of what types of ORFs we define as “non-canonical”. Yet, while analyses of more cell lines and tissues will certainly uncover additional non-canonical ORFs, there can be variable non-canonical ORF identifications even within analyses of the same cell line. Such variability reflects the equal – perhaps foremost – contribution of different analytical methods for non-canonical ORFs in the estimation of their prevalence.

#### The number of non-canonical ORFs

Most Ribo-seq studies focusing on non-canonical ORFs report detection of several thousand ORFs, typically between 2,000 and 8,000 (9, 11, 15, 16, 18–21, 51, 61, 114). Interestingly, this range seems relatively stable when comparing studies that employ only a few cell lines and broader analyses looking across many different human tissue types. To consolidate these findings, we have recently participated in an international consortium to aggregate 7,264 high-confidence non-canonical ORFs and provided formalized annotations for them within the GENCODE gene annotation database (16). This GENCODE set demonstrates substantial overlap in the identification of certain types of ORFs, such as upstream ORFs (uORFs), across diverse datasets such as pancreatic progenitors, heart and stem cells, suggesting that perhaps the diversity of several ORF types may not be dramatically larger with the inclusion of more tissue types. In support of this, Ribo-seq profiling of five human tissue types and 6 primary human cell types similarly reported 7,767 ORFs in total (15). When subsetting this dataset for consistency with the inclusion criteria for the GENCODE catalog (i.e., removing ORFs below 16 amino acids in size, as well as ORFs without an AUG start codon), 2,475 out of 7,767 ORFs remained, of which 1,702 (± 70%) were represented in the GENCODE catalog as well (**Supplementary Tables 1-4**).

While these studies have measured and determined non-canonical ORF translation directly from Ribo-seq data, there are many other databases that have aggregated larger numbers of ORFs from a variety of sources, including both Ribo-seq and *in silico* predictions. Among these, smProt (n = 327,995 human ORFs (115)), sORFs.org (n = 4,377,422 ORFs across humans, mouse and fruit flies (116)), RPFdb (117, 118) and smORFunction (n = 617,462 human ORFs (119)) have compiled reported or putative non-canonical ORFs. Notably, OpenProt (120, 121) has two aspects to their database workflow: one that collates all predicted ORFs (n = 488,956) and a second that proposes 33,836 translated ORFs identified by a re-analysis of over a hundred Ribo-seq datasets with the PRICE pipeline (67). When considering studies that have generated Ribo-seq datasets to measure non-canonical ORF translation, there are also several efforts that have proffered exceptionally large numbers of directly detected ORFs – specifically the nuORFdb (34) by Ouspenskaia *et al.* and the Human Brain Translatome Database (122) by Duffy *et al.*, which propose numbers of >230,000 and >75,000 ORFs, respectively.

#### Why is there such discordance in the number of non-canonical ORFs across databases?

The interpretation of such dramatically different accounts of non-canonical ORF abundance remains a challenge. Indeed, given that there are currently only ∼60,000 Ensembl genes (including 19,827 protein-coding genes, 18,886 lncRNAs, 4,864 small ncRNAs, 15,241 pseudogenes, and 2,221 other RNAs in Ensembl v109.38), colossal datasets with >200,000 ORFs may be interpreted to suggest that every gene has upwards of 4 distinct ORFs. In practice, these large datasets may include isoform variants (e.g. N-terminal extensions, C-terminal extensions and intron retentions) that are not part of the reference proteome, and thus the number of non-canonical ORFs may be larger in some databases due to differences in how these isoforms are categorized.

While sample and data quality likely contribute to the variability in the numbers of non-canonical ORFs in some catalogs, differences in Ribo-seq data analysis also account for much variation in prospective non-canonical ORFs. For example: biologically, there is some amount of stochastic or pervasive translation across all RNAs, which may relate to leaky ribosomal scanning (123–125) or transient interactions between ribosomes and RNAs as the ribosomes locate coding sequences or RNAs accomplish proper folding (126, 127). Yet, the manner in which computational pipelines process Ribo-seq data results in ORF calls that may be more or less stringent (**Figure 2B**), resulting in different proportions of false-positive (stochastic) and false-negative (e.g., sample-specific) ORF calls (60, 128, 129). For example, RibORF (19), which uses a support vector machine and recommends a fixed cut-off score of 0.7, has been shown to produce the highest numbers of ORF calls of any tested algorithm in a recent benchmarking study (130). To confirm these differences directly, we have re-analyzed published high-quality Ribo-seq data for six biological replicates of pancreatic progenitor cells differentiated from human embryonic stem cells (11) using four common ORF detection pipelines (ORFquant (10), PRICE (67), Ribo-TISH (66), and Ribotricer (69)), observing substantial variability in the number of ORFs called (∼10-fold difference from ∼50,000 to ∼500,000), the types of ORFs called, the length of the called ORFs, and the reproducibility with which ORFs could be detected across all six replicates (**Figure 3** and **Methods**).

**Figure 3:**
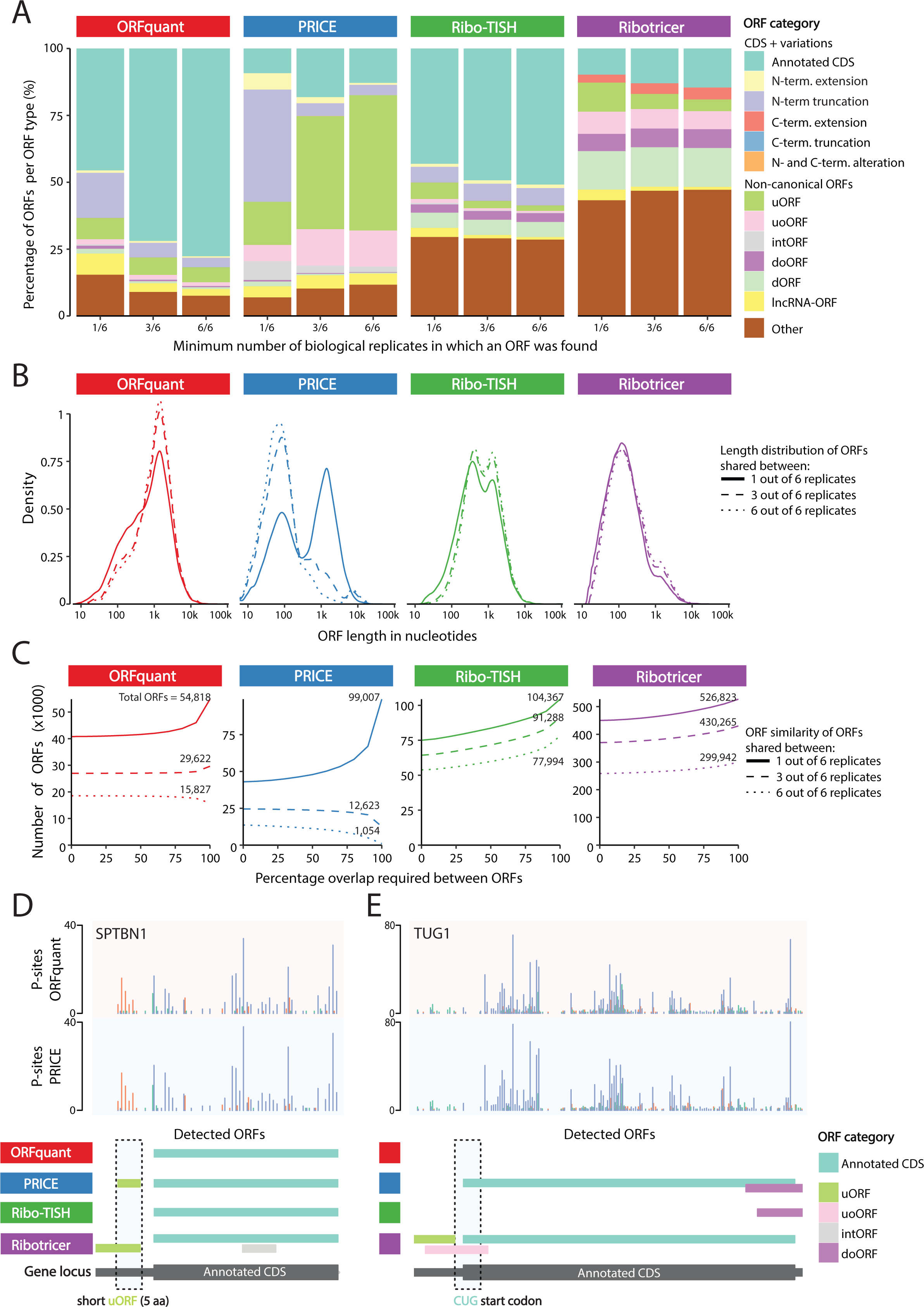
ORF callers have different specialties and variable performance. A) Stacked bar plot displaying all detected ORF categories per ORF caller. For each, the percentage of unique ORFs shared between at least one, three, or six replicates is shown. Please note that these are relative contributions to the total number of ORFs. The absolute numbers of ORF identifications can be inferred from Figure 3C. B) Density plots displaying the distribution of ORF lengths in nucleotides (excluding the stop codon) for unique ORFs shared between at least one, three, or six replicates. C) Line graphs showing the numbers of unique ORFs detected by each tool shared between at least one, three or six replicates. The x-axis denotes the percentage of overlap used to consider two ORFs being similar or not, with 100% overlap meaning that the detected ORF was fully identical between [x] number of replicates. Please note that the total numbers of ORFs detected per algorithm (y-axis) can differ by an order of magnitude. These numbers are given for each line, with numbers reflecting the total ORFs with 100% similarity between replicates (i.e., the end of each curve). D) Genomic view of a short upstream ORF (uORF) in the STPBN1 gene indicating that ORF callers have variable affinity for certain types of ORFs. The top two tracks show the ribosomal P-site positions derived from the sequenced ribosome footprints, as processed independently from the sequencing data by the deterministic ORF caller ORFquant (top; red shading) and the probabilistic ORF caller PRICE (bottom; blue shading). The differently colored P-site bars indicate different reading frames (0, +1, +2) on the same transcript, with bars in the same color indicating a shared in-frame codon movement by the ribosome. For this visualization, newly found ORF variations of the annotated CDS that could be assigned to predicted non-coding RNA isoforms (e.g., transcript biotype: “processed_transcript”), but matched the CDS of SPTBN1 is not displayed. E) Genomic view of a near-cognate start codon ORF in TUG1. Image and track details as in (E) above.

There may be specific reasons for the different performance characteristics of each algorithm. For example, the lower stringency of RibORF may be due to the fact that this pipeline considers uniformity of read coverage across the ORF, while Ribo-seq is known to have a 5’ bias to read coverage. Therefore, RibORF may excessively promote intORFs and doORFs since the 5’ ends of these ORFs overlap annotated CDSs, which typically have higher read coverage independent of a periodic footprint signal that matches the correct reading frame. This is evident in nuORFdb (34) and the Human Brain Translatome Database (122): when analyzing the fraction of ORFs with an AUG-start resulting in an ORF >=16 amino acids, doORFs and intORFs are 173-fold and 18-fold (respectively) higher in abundance compared to other major datasets (**Figure 4, Supplementary Tables 5-8**). By contrast, uORFs are only 3 times more abundant (**Figure 4**).

**Figure 4:**
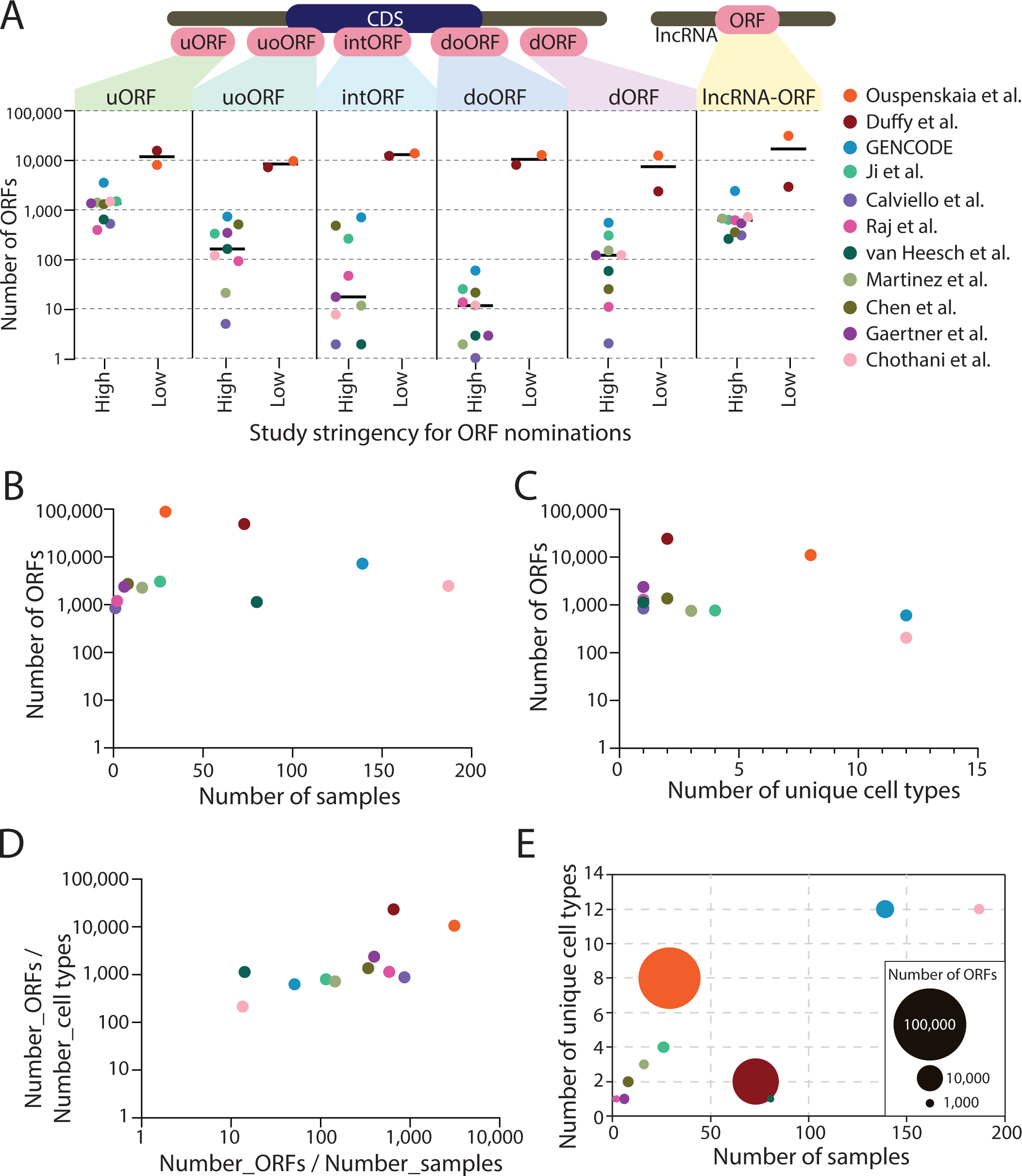
An analysis of major non-canonical ORF databases. A) Here, each dot reflects a dataset, and the Y axis uses a log-10 scale to show the number of ORFs included that are >=16 amino acids long and contain an AUG start codon. The GENCODE catalog reflects the summation of the Ji et al. (19), Calviello et al. (61), Raj et al. (20), van Heesch et al. (9), Martinez et al. (21), Chen et al. (18) and Gaertner et al. (11) datasets as described in (16). B) The number of ORFs per dataset compared to the number of samples profiled by Ribo-seq. C) The number of ORFs per dataset compared to the number of unique cell types profiled by Ribo-seq D) The ratio of the number of ORFs per cell type compared to the number of ORFs per number of samples for each dataset. E) A bubble plot integrating the number of samples, number of different cell or tissue types, and the number of non-canonical ORFs found in each dataset.

It is also true that different computational pipelines may have different capacity to identify certain classes of non-canonical ORFs. For example, the deterministic multitaper-based statistical inference of significant periodic signal within predicted ORFs as performed by Ribotaper (61) and ORFquant (10) provides high-confidence detection of ORFs with an AUG start codon, but have not, to date, been optimized for non-AUG ORFs. In contrast, the probabilistic algorithm employed by PRICE (67) has enhanced ability to identify very short ORFs and non-AUG ORFs absent from other ORF callers (**Figure 3B+E**). Yet, when there are neighboring putative initiation codons (e.g. CUG and AUG), PRICE will generate larger numbers of putative ORFs that might require manual curation or further filtering. In addition, since annotated CDSs have generally more abundant Ribo-seq read coverage, low-abundance out-of-frame reads may be more readily interpreted as an intORF with a non-AUG start codon by PRICE, whereas other ORF callers are less likely to consider these reads as sufficient evidence for a translated ORF. Thus, when applied to biological replicates of the same sample, PRICE produces the least consistent ORF calls compared to other pipelines, independent of initiation codon variability (**Figure 3A-C**) (130). nuORFdb (34) and OpenProt (121) both employ PRICE in their analysis pipelines. It is important to note, however, that the specific research question being pursued should inform the types of ORF callers used: indeed, deterministic algorithms such as RiboTaper or ORFquant may miss intORFs or overlapping ORFs identified by PRICE because of the difficulty in resolving mixed periodicity signals of overlapping reading frames (**Figure 3A**).

In summary, depending on the type of ORF one aims to find and the desired inclusiveness of ORFs one aims to output, one ORF caller might be better suited than another. Certain ORF callers outperform others in detecting specific ORF categories such as intORFs (**Figure 3A**), very small ORFs (**Figure 3B+D**), or near cognate start codons (**Figure 3E**), whereas others handle exon-exon junctions and longer ORFs better and/or provide better replicate behavior. These differences then lend to substantially different results when producing non-canonical ORF catalogs (**Figure 4**).

#### Detection of translational start sites

Determining the translational start site of an open reading frame remains a nuanced problem. While conventionally proteins have been annotated with AUG start sites, exceptions to this rule have long been known (131, 132), and non-canonical ORFs are more likely to employ non-AUG start sites (124, 133). In a typical Ribo-seq experiment, identification of translational start sites from Ribo-seq data is inferred based on two factors: sequencing coverage and the intrinsic restrictions of the computational pipeline (e.g. some algorithms only consider AUG start codons, as discussed above). Yet, independent of the computational pipeline, there may be gaps in the sequencing coverage that lead to mis-identification of the main translational initiation site (**Figure 2C**). For experiments with cultured cells, use of small molecules that block ribosome elongation, such as homoharringtonine (134) or lactimidomycin (135), enables ribosome accumulation on translational initiation sites, which enables more precise determination of the start codon. Due to the difficulty in identifying non-canonical ORF start sites and the variability in computational approaches to start codon recognition (e.g., **Figure 3E**), use of homoharringtonine or lactimidomycin with cultured cells is highly recommended. In frozen tissue samples, these compounds are no longer effective.

### How to select an ORF sequence database for MS data analysis

Given the wide differences between the different databases for Ribo-seq ORFs, one central question is how to use these databases, or which to use for any specific analysis? Because the size of the ORF output in a given database can vary enormously, users should base their decision on what scientific question they intend to pursue and evaluate carefully the suitability of the input Ribo-seq data quality as well as the stringency with which ORF calling was performed. In general, high stringency databases provide high confidence Ribo-seq ORF detections, and thus peptides found mapping to these ORFs are more likely to reflect a true positive result. While these databases reduce false positives, it is at the expense of comprehensiveness, as the existing high stringency databases will yield more false negatives in the MS analysis. Low stringency databases provide a much larger set of Ribo-seq ORFs, but will yield more false positives – due to the lack of support from another orthogonal technique. If the ORFs are accompanied by Ribo-seq quality metrics, it may be tractable to estimate the proportion of false positives, and re-filter the ORFs to suit one’s own purposes. These databases will provide a larger candidate search space for peptide alignment and may enable detection of true positive ORFs not present in the high stringency databases. Yet as described earlier, due to the concern for false positive nominations, ORFs detected by mass spectrometry searches should be closely inspected to verify integrity of both ORF call and peptide identification, as there will likely be cases of false-positive ORFs being supported by false-positive peptides. Ultimately, certain scientific questions may lend themselves to certain databases: for example, analyses of alternative N-terminal CDS extensions often emphasize non-AUG start sites (24), which may benefit from a Ribo-seq analysis that employs the PRICE algorithm. Research efforts aimed to identify a maximal space of potential translation events may also favor a lower stringency database, with the caveat that any individual result should receive additional scrutiny. Alternatively, if the goal is to characterize a high-confidence unannotated microprotein, a high stringency database may be more desirable. Likewise, for reference annotation purposes and functional studies we prefer more stringent workflows that yield reproducible ORF calls across samples (no false positives).

### Are non-canonical ORFs proteins?

The term “protein” is conventionally used to refer to an amino acid sequence that produces a molecular structure that plays an intrinsic cellular role in maintaining normal cell biology. While some proteins may be unstable and rapidly degraded under certain conditions (e.g., beta-catenin), most proteins participate in cell biology when present in a stable form. Also, almost all annotated proteins show evidence of evolutionary conservation, structural folding and domain architecture, and frequently also protein-protein interactions and/or interactions with nucleic acids.

According to this understanding of the term “protein”, it could be inferred that the vast majority of non-canonical ORFs do not encode proteins on the basis that they lack these characteristics. However, we see two additional considerations. First, it may be incorrect to assume that a protein that exists in the cell – even one that is detectable by mass spectrometry – is therefore a functional molecule. It could be that the proteome contains a certain amount of non-functional translational “noise”. While it is difficult to prove the extent to which such translation occurs in normal cells, evidence from cancer cells shows abundant dysregulation of translation, exemplified by “aberrant” non-canonical proteins that lack evidence for function under normal physiological conditions (34, 35) as well as out-of-frame peptide byproducts of oncogene activity (136).

Second, the classical definition of protein “function” invokes the protein’s role in cellular processes that have been derived over time through evolution, which has been summarized as the maxim that “conservation = function”. This maxim has been central – but not universally required – for gene annotation projects, and the only canonical proteins currently within GENCODE that can be inferred to have evolved *de novo* in human or higher primates were initially detected in cancer cells (e.g. MYEOV (137) and HMHB1 (138)). Even so, evidence for the existence and function of *de novo* proteins under normal physiological conditions is accumulating (57, 139–141). Nonetheless, it remains true that most non-canonical ORFs display much higher rates of intrinsic disorder, fewer structural features, and lack amino acid constraint across evolution (17, 18, 139, 140, 142–148). While these features may be observed in diverse annotated proteins (e.g. intrinsically disordered regions of a given protein), their presence is predominant in non-canonical ORFs.

#### The interpretation of peptide-level evidence of Ribo-seq ORFs

How, then, should one interpret the peptide-level evidence for some non-canonical ORFs? High-quality whole proteome LC-MS/MS PSMs that survive rigorous manual inspection are strong evidence of true translation of a non-canonical ORF. With adequate evidence, therefore, whole proteome PSMs supporting non-canonical ORFs do indicate the possible existence of a translated protein, and these cases may reasonably be considered to be part of the cell proteome, similar to any other proteins.

When considering the larger number of non-canonical ORFs with peptide-level evidence in HLA immunopeptidomics but not shotgun LC-MS/MS (18, 34, 36, 38, 149), firm conclusions are more difficult to draw. These non-canonical ORFs cannot be said to generate a true protein based on immunopeptidomics alone, considering that the HLA system is expected to present peptides resulting from translation products that are unstable and rapidly degraded, alongside those derived from canonical proteins. Yet, detection of an HLA-presented peptide does verify RNA translation in these cases, which distinguishes them from the majority of Ribo-seq-detected non-canonical ORFs that are detected in neither shotgun LC-MS/MS nor immunopeptidomics experiments. Therefore, these non-canonical ORFs can at least be said to be confirmed as both translated and presented by the HLA, as opposed to an artifact of the Ribo-seq protocol.

A related question is how to interpret PSMs matching non-canonical ORFs that are not detected by Ribo-seq, when the same sample is interrogated using both technologies. Because the sensitivity of Ribo-seq is generally higher than MS-based methods, and because Ribo-seq provides nucleotide-level precision for genome mapping, there are three possibilities here: first, these peptides may be false-positive identifications, second, the Ribo-seq data exhibit a false-negative, or third, they may be derived from another source not included in the search space (ex: aberrant splicing). None of these hypotheses has been rigorously evaluated at this time. One challenge is that many proteomics and immunopeptidomics experiments do not currently generate matched Ribo-seq data for their samples, and thus it cannot be directly known if Ribo-seq supports translation of that ORF. When considering unmatched analyses, it is also noted that, at present, proteomics and immunopeptidomics datasets cover a broader range of tissue and cell types than Ribo-seq datasets.

### A proposed framework to classify the translation of non-canonical ORFs

Given the expanding volume of research on non-canonical ORFs, a shared vocabulary for the interpretation of their detection is a critical need in the genomics, translatomics, proteomics, and immunopeptidomics communities. Notably, there has been no formalized initiative to annotate non-canonical ORFs as protein-coding genes by major genome databases, although recent collaborative work has raised this point as a topic of interest (16). Historically, protein-coding genes have been annotated one-by-one in a manual process of careful data inspection, which may or may not have included protein-level evidence. At this time, non-canonical ORFs detected by tryptic whole proteome data would potentially be eligible for manual annotation as protein-coding genes. Yet, given the paucity of non-canonical ORFs in tryptic whole proteome data and their much greater abundance in HLA immunopeptidomic datasets, there is uncertainty about whether most non-canonical ORFs produce proteins in the classical sense, and whether immunopeptidomic evidence is equivalent to tryptic whole proteome data for the purposes of protein annotation.

We advocate both a cautious but open-minded approach to non-canonical ORF classification, summarized in **Table 2**. Notably, although most annotated proteins show evidence of amino acid constraint across species and most non-canonical ORFs do not, it is also unquestionably true that at least some proteins are lineage- or species-specific. Thus, we propose that *de novo* translations should be considered for annotation as protein-coding. However, recognizing that evolutionary analysis is a core part of gene annotation workflows in projects like GENCODE, we have not included conservation or constraint metrics as part of this proposed framework. The framework itself is oriented toward harmonizing subsequent dataset generation and analysis. In practice it might be applied to classifying published datasets, and it is intended as a helpful tool for candidate prioritization rather than a guarantee that certain ORFs will be annotated by a genome database. We stress that researchers looking to move forward with potential annotation of a protein encoded by a non-canonical ORF should be able to provide the raw LC-MS/MS spectra for review.

**Table 2.** A proposed framework to standardize levels of evidence of non-canonical ORFs.

| Tier | Required Supporting Evidence | Standardized Outcome |
| --- | --- | --- |
| *Tier 1A* | Whole proteome LC-MS/MS ($\geq 2$ peptides according to HUPO HPP criteria)<br>Ribo-seq* | *"Protein candidate"*<br>Consider discussing research findings with genome annotation databases for possible annotation. |
| *Tier 1B* | HLA immunopeptidomics MS ($\geq 2$ observations; multiple high confidence peptides from multiple distinct samples)<br>Ribo-seq* | *"Presented"* |
| *Tier 2A* | Whole proteome LC-MS/MS ($\geq 2$ peptides not satisfying HUPO HPP spacing criteria)<br>Whole proteome LC-MS/MS (1 peptide)<br>Ribo-seq* | *"Detected"* |
| *Tier 2B* | HLA immunopeptidomics MS (1 observation)<br>Ribo-seq* | *"Detected"* |
| *Tier 3* | Any HLA immunopeptidomics or whole proteome LC-MS/MS evidence without Ribo-seq* evidence | *"Putative", consider alternative sources.* |
| *Tier 4* | Ribo-seq* evidence without any proteomic evidence | *"Ribo-seq ORF"* |
| *Tier 5* | *In silico* prediction of an ORF on an expressed transcript without any Ribo-seq* or proteomic evidence | *"Predicted"* |

\*from credible Ribo-seq data with quality metrics meeting the guidelines suggested in this manuscript. Ribo-seq need not be performed on aliquots of the same samples analyzed by proteomics.

Our framework proposes these definitions for specific terminology:

- *“Protein candidate”:* a **Tier 1A** non-canonical ORF can be regarded as translated into a protein candidate if it satisfies current HUPO HPP guidelines for the detection of >= 2 uniquely mapping peptides, as well as having evidence of translation by Ribo-seq. Such candidates would be prioritized for further manual review by annotation groups.
- *“Presented”*: A presented non-canonical ORF (**Tier 1B**) is one with multiple lines of evidence for its translation and presentation on HLA molecules. These ORFs are detected with multiple high confidence peptides from multiple distinct samples for HLA immunopeptidomics data, as well as having evidence of translation by Ribo-seq.
- *“Detected”*: A detected non-canonical ORF is one with evidence of translation by Ribo-seq as well as evidence of protein production by either (**Tier 2A**) shotgun LC-MS/MS (1 peptide or >1 peptide not satisfying HUPO HPP guidelines for their spacing), or evidence of protein production by HLA immunopeptidomics with a single PSMs (**Tier 2B**).
- *“Putative”*: A putative non-canonical ORF (**Tier 3**) is one with evidence of translation with LC-MS/MS or HLA immunopeptidomics data, but no evidence of translation in Ribo-seq data. This discrepancy may alert to the possibility of false-positive MS identifications, or false-negative absence in Ribo-seq, and therefore requires more investigation.
- *“Ribo-seq ORF”*: A non-canonical ORF that is only detected in Ribo-seq data but not elsewhere is considered a “Ribo-seq ORF” (**Tier 4**). These are likely to be the majority of cases. The number of these ORF nominations may be variable based on the stringency of the Ribo-seq analysis and/or the quality of the input data.
- *“Predicted”*: A predicted non-canonical ORF (**Tier 5**) is one that is computationally predicted *in silico* on an expressed RNA transcript, but without current evidence in Ribo-seq or MS datasets.

## Limitations

With this work, we have endeavored to clarify how Ribo-seq can be used for non-canonical ORF research. Yet, our focus has several important limitations. First, the vast majority of – but not all – translated peptides can be traced back to an RNA sequence. There may be peptides that derive from amino acid splicing within the proteasome during protein degradation (150), which would not be detectable in Ribo-seq data. Second, there are also well-established protein-coding sequences that are difficult to resolve with Ribo-seq and do not have optimized computational methods for their quantification. For example, translated pseudogenes, retroviruses, retrotransposons, and paralogous protein-coding genes may have high sequence homology that precludes unique mapping of the short ∼30 bp reads from a Ribo-seq experiment, although multi-mapping reads will provide evidence of translation. These cases are not discussed here. Lastly, each individual’s genome (and particularly each cancer’s genome) has a unique range of germline or somatic single nucleotide variants that will impact the proteome: in this article, we have not addressed the importance of generating personalized reference genomes and proteomes for the analysis of microproteins and non-canonical ORFs.

## Conclusions

The widespread description of non-canonical ORFs has sparked a paradigm shift in the perception of both the human genome and the proteome. Yet, as a field still in its infancy, this area of investigation is plagued by a lack of standardization, which may lead to imprecise analyses, ultimately leading to self-injurious confusion. While the proportion of non-canonical ORFs that encode a functional protein remains to be seen, a large fraction of them can be verified as translated by both MS-based and Ribo-seq-based approaches. A central effort for the research community is now to build reputable databases and analysis pipelines to ensure rigor in this quickly-expanding – and highly exciting – field. Here, we have considered the technologies used to detect non-canonical ORFs and attempted to provide a framework for categorizing differing levels of evidence for them. Our work aims to coalesce the research community around a common terminology and shared set of database resources for non-canonical ORFs. Ultimately, we believe that the study of non-canonical ORFs, if pursued with proper precision, will prove invaluable to the global community of biomedical researchers.

## Abbreviations

HLA: human leukocyte antigen
MHC: major histocompatibility complex
CDS: coding sequence
LC-MS/MS: liquid chromatography with tandem mass spectrometry
MS: mass spectrometry

## Acknowledgements

J.R.P. acknowledges funding from the National Institutes of Health/National Cancer Institute (K08-CA263552-01A1), the Alex’s Lemonade Stand Foundation Young Investigator Award (#21-23983), the St. Baldrick’s Foundation Scholar Award (#931638), the Musella Foundation for Brain Tumor Research, the DIPG/DMG Research Funding Alliance, and a Collaborative Pediatric Cancer Research Awards Program/Kids Join the Fight award (#22FN23). E.W.D. acknowledges funding from National Institutes of Health grant R01 GM087221 and National Science Foundation grant DBI-1933311. S.v.H. acknowledges funding from Fonds Cancers, Stichting Reggeborgh, Stichting Bergh in het Zadel, and Stichting Villa Joep. J.G.A and K.R.C were supported in part by grants P01CA206978 from the NIH, U24CA270823, U01CA271402 and U24CA271075 from National Cancer Institute (NCI) Clinical Proteomic Tumor Analysis Consortium program and from the Dr. Miriam and Sheldon G. Adelson Medical Research Foundation. J.M.M. is supported by the Wellcome Trust (grant number 108749/Z/15/Z), the National Human Genome Research Institute (NHGRI) of the US National Institutes of Health (NIH) under award number 2U41HG007234, and the European Molecular Biology Laboratory (EMBL). The content is solely the responsibility of the authors and does not necessarily represent the official views of the National Institutes of Health. Ensembl is a registered trademark of EMBL.

## Author Contributions

Conceptualization: E.W.D., J.R.P., S.v.H., J.R.-O., J.M.M., M.B.-S., J.G.A., K.R.C.

Methodology, E.W.D., J.R.P., S.v.H., J.R.-O., J.M.M., M.B.-S., J.G.A., K.R.C., L.W.K.

Formal analysis, J.R.P., L.W.K

Data curation, J.R.P., J.R.-O., L.W.K.

Writing - E.W.D., J.R.P., S.v.H., J.R.-O., J.M.M., M.B.-S., J.G.A., K.R.C.

Writing - review & editing, E.W.D., J.R.P., S.v.H., J.R.-O., J.M.M., M.B.-S., J.G.A., K.R.C., L.W.K.

Visualization, J.R.P., L.W.K.

Supervision, E.W.D., J.R.P., S.v.H., J.R.-O., J.M.M., M.B.-S., J.G.A., K.R.C.

Funding acquisition, E.W.D., J.R.P., S.v.H.

## Inclusion and diversity

We support inclusive, diverse and equitable research.

## Methods

### Benchmarking and comparing ORF caller performance on replicate Ribo-seq datasets

#### Ribo-seq data processing and mapping

Ribosome profiling data of late pancreatic progenitor cells obtained from six independent differentiations of H1 human embryonic stem cells (11) were collected from the GEO database (GSE144682). For all analyses, the Ensembl primary DNA assembly (GRCh38) and the Ensembl human reference transcriptome (Ensembl v102) were used as reference. Quality control and trimming of the Ribo-seq reads was done using Trim Galore 0.6.6 with the options ‘--length 25’, and ‘-- trim-n’ (151). Next, contaminant RNA and DNA were removed using Bowtie2 2.4.2 by aligning reads to a contaminant file using the default options of Bowtie2 (152). The contaminant-depleted reads were aligned using STAR with the options ‘--twopassMode Basic’, ‘--outFilterMismatchNmax 2’, ‘-- outFilterMultimapNmax 20’, ‘--limitOutSJcollapsed 10000000’, ‘--alignSJoverhangMin 1000’, and ‘-- outSAMattributes All’ (153). For PRICE, the option ‘--alignEndsType EndToEnd’ was set as well. Also, the individual bamfiles were filtered using SAMtools 1.12 to exclude reads with a mapping quality lower than 5 (154).

#### ORF calling with ORFquant

The function RiboseQC_analysis from RiboseQC 1.1 was run in R 4.1.2 with the options ‘read_subset’ and ‘fast_mode’ set to false (155). The output was used by the function run_ORFquant from ORFquant 1.02 in R with the default options (10). ORF calling with PRICE: Before using PRICE, a reference genome was created with the IndexGenome function of the Gedi framework 1.0.2. After the creation of the reference genome, PRICE 1.0.3b was run (67). A filtered list of ORFs detected by PRICE and a list of P-sites (called activity values by PRICE) were extracted from the outputted ‘orfs.cit’ files using the Gedi Nashorn and ViewCIT functions respectively. Because the start codon prediction is a separate step in the PRICE program, ORF coordinates from both before and after start codon prediction were available. We used the coordinates after start codon prediction. PRICE can also be run in a multi sample mode by providing a text file with the bam file locations as input. This mode favours ORFs that occur in all samples during the ORF calling process and would likely enhance the reproducibility of ORF calls between replicates. To keep all ORF callers comparable we did not use this mode. ORF calling with Ribo-TISH: From Ribo-TISH 0.2.7, the predict function was used to infer ORFs with the option ‘--longest’ set (66). The output file contained only the genomic start and end coordinates and the transcript id of each ORF. The reference GTF was used to determine the exons within each ORF. ORF calling with Ribotricer: The Ribotricer 1.3.3 function prepare_orfs was first used with the options ‘--longest’, ‘-- min_orf_length 9’ (69). The option ‘--start_codons’ was set to include all near cognate start codons with one base difference compared to ATG. Afterwards, the function “detect_orfs” was used with the option ‘--phase_score_cutoff 0.440’.

#### Comparing ORF callers

ORF calls were compared between algorithms for the types of ORF categories that were found, in how many replicates they were independently discovered, how ORF differed in length, and how reproducible and similar their detection was based on e.g., the percentage of ORF sequence overlap between replicate ORF calls. Before the analyses, data were converted to GRangesList objects in R with stop codons included in the coordinates. ORF categories were determined by comparing the start and end coordinates, and the transcript id of each ORF with the coding sequences in the ‘gtf.rannot’ object created by the ORFquant function ‘prepare_annotation_files’. ORFs were compared by their overlap, with different thresholds set for the required percentage of overlap. Two ORFs were considered to be similar if the exons of one ORF were fully contained within the exons of a second ORF, both codons had the same stop codon, and the first ORF covered at least the required percentage of overlap of the length of the second ORF. These overlap relations were recursive, such that a parent ORF could be the child of another ORF, and all three would be counted as one unique ORF.

#### Code availability

All code used for these analyses as well as data visualization are available at https://bitbucket.org/vanHeeschLab/orfcaller_comparison.

### Comparison of published Ribo-seq datasets

We used publicly-available datasets from GENCODE (16), Chothani et al. (15), Ouspenskaia et al. (34), and Duffy et al. (122) for comparisons of published reports of non-canonical ORFs that might encode microproteins. The GENCODE dataset itself is a metaanalysis of data from Ji et al. (19), Calviello et al. (61), Raj et al. (20), van Heesch et al. (9), Martinez et al. (21), Chen et al. (18), and Gaertner et al. (11); datasets employed are listed in **Supplementary Table 1**. Source data for these datasets is listed in **Supplementary Table 2**. To facilitate comparisons between studies, we extracted only non-canonical ORFs with a length of >=16 amino acids and had an AUG start codon. For ORFs using a non-AUG start site, the first internal AUG start codon was identified and the amino acid sequence starting with that internal AUG was included for analysis if the resulting ORF was >=16 amino acids long. ORFs were then analyzed for their replication across primary datasets. Since the GENCODE list represents a metaanalysis of other individual datasets, presence of an ORF in the GENCODE list was not used as part of the analysis for ORF replication across primary datasets. Next, ORF calls were associated with one of the following six categories: lncRNA-ORF, upstream ORF, upstream overlapping ORF, internal ORF, downstream overlapping ORF, or downstream ORF, according to schema by Mudge et al. (16). Duffy et al. used the nomenclature “external” for doORF, and these ORFs were reclassified as doORF for this analysis; they used “internal” for intORFs, which were reclassified as intORFs for this analysis. For lncRNAs, Duffy et al. used the term “non-coding” which included the biotypes “noncoding”, “lncRNA”, “antisense_RNA”, “misc_RNA”, “TEC” and “processed_transcript”, which were included as part of the lncRNA-ORF designation for this study. For Ouspenskaia et al., we analyzed ORFs according to the authors’ designation of ORF “plotType”, reflecting their final classification. Ouspenskaia et al. used the term “3’ dORF” for dORF, “3’ overlap dORF” for doORF, “5’ overlap uORF” of uoORF, “5’ uORF” for uORF, “lncRNA” for lncRNA-ORF, and “out-of-frame” for intORF. Chothani et al. reported final ORF types of “dORF”, “doORF”, “ncORF”, “overlap_uORF”, “intORF” and “uORF”. For Chothani et al., Duffy et al. and Ouspenskaia et al., ORFs that had a final classification of pseudogene were excluded from this analysis; however, these datasets variably re-classified some ORFs on pseudogene transcript biotypes as non-coding or lncRNA, and we did not re-filter these ORFs beyond the original reclassifications provided by the authors. ORFs switch a classification corresponding to a small RNA, tRNA or rRNA species, such as “rRNA”, “snoRNA”, “tRNA”, “snRNA” or “miRNA”, were excluded from this analysis. The number of cell types and/or tissue types for analyses of each ORF dataset was extracted from the source publication.

## Supplementary Materials

### List of Supplementary Tables

Supplementary Table 1: Description of aggregated data

Supplementary Table 2: A description of ORF primary datasets and data sources

Supplementary Table 3: Phase I Ribo-seq ORFs detected in only 1 of 8 studies

Supplementary Table 4: A metaanalysis of non-canonical ORFs present in 8 different high-stringency datasets

Supplementary Table 5: ORFs included for this analysis from the Duffy et al. dataset.

Supplementary Table 6: ORFs included for this analysis from the Ouspenskaia et al. dataset.

Supplementary Table 7: The number of ORFs for comparative analysis across studies

Supplementary Table 8: The number of ORFs per sample and per cell/tissue type according to each study

